# Reconceptualizing programmed transcriptional slippage in RNA viruses

**DOI:** 10.1101/2024.02.05.578984

**Authors:** Adrian A. Valli, María Luisa Domingo-Calap, Alfonso González de Prádena, Juan Antonio García, Hongguang Cui, Cécile Desbiez, Juan José Lopez-Moya

**Affiliations:** Centro Nacional de Biotecnología (CNB-CSIC), 28049 Madrid, Spain; Center for Research in Agricultural Genomics (CRAG-CSIC-IRTA-UAB-UB), Campus UAB, 08193 Bellaterra, Spain; Evolving Therapeutics SL., Parc Científic de la Universitat de València, 46980 Paterna, Spain; Key Laboratory of Green Prevention and Control of Tropical Plant Diseases and Pests, Ministry of Education and College of Plant Protection, Hainan University, Haikou, Hainan, China; INRAE, UR0407 Plant Pathology, F-84143 Montfavet, France

## Abstract

RNA viruses have evolved sophisticated strategies to exploit the limited encoded information within their typically compact genomes. One of such, named programmed transcriptional slippage (PTS), is defined by the insertion of an additional A at A_n_ motifs (n ≥ 6) of newly synthetized viral transcripts to get access to overlapping open reading frames (ORFs). Although key proteins from Ebolavirus and potyvirids (members of the *Potyviridae* family) are expressed via PTS, available information about this phenomenon is very scarce. Here, by using diverse experimental approaches and a collection of plant/virus combinations, we discover cases in which PTS does not fit with its current definition. In summary, we observe (i) high rate of single nucleotide deletions at slippage motifs, (ii) overlapping ORFs acceded by slippage at an U_8_ stretch, and (iii) significant changes in slippage rates induced by factors not related to cognate viruses. Moreover, a survey of full-genome sequences from potyvirids shows a widespread occurrence of species-specific A_n_/U_n_ (n ≥ 6) motifs. Even though many of them, but not all, lead to the production of truncated proteins rather than access to overlapping ORFs, these results suggest that slippage motifs appear more frequently than expected and play relevant roles during virus evolution. In conclusion, our data prompt to broaden PTS definition in RNA viruses. Considering the potential of this phenomenon to expand the viral proteome by acceding to overlapping ORFs and/or producing truncated proteins, a revaluation of PTS significance during infections of RNA viruses is required.

**IMPORTANCE:** Programmed transcriptional slippage (PTS) is used by RNA viruses as another strategy to maximise the coding information in their genomes. This phenomenon is based on a peculiar feature of viral replicases: they insert an untemplated A in An motifs (n ≥ 6) in a small fraction of newly synthesised viral RNAs. As a consequence, ribosomes can get access to overlapping open reading frames (ORFs) when translating those particular transcripts. Here, using plant-infecting RNA viruses as models, we discover cases challenging the previously stablished definition of viral PTS, prompting us to reconsider and redefine this expression strategy. An interesting conclusion from our study is that PTS might be more relevant during RNA virus evolution and infection processes than previously assumed.

## INTRODUCTION

RNA viruses possess small genomes, resulting in limited coding capacity. Possibly in response to this constraint, they have evolved diverse and intricate gene expression strategies. One of such strategy involves the generation of *de novo* proteins via overprinting -a process wherein specific nucleotide substitutions in an existing gene prompt the expression of a novel protein from an alternative open reading frame (ORF) (2). Overprinted genes can be accessed through either transcriptional or translational mechanisms. In the former, the alternative RNA is transcribed from internal promoters (3) or through programmed transcriptional slippage (PTS) (4, 5). In the later, the protein is produced via translation from alternative start codons, or by ribosomal read-throughs and frame-shifts (5, 6). Of these mechanisms, PTS is the least characterized, with only few well-defined cases described across unrelated taxonomic groups. Three examples are comprised by the expression of the large glycoprotein (GP) from Ebolavirus (7) and both P3N-PIPO and P1N-PISPO from potyviruses (members of the *Potyvirus* genus, *Potyviridae* family) (8–10). Despite the key relevance of these proteins for viral infection, the precise molecular mechanism by which PTS takes place is still a matter of study (11, 12). So far, it is known that viral RNA-dependent RNA polymerases (RdRPs) slip over homopolymeric runs of A_n_ (n ≥ 6) when copying the viral genome - or U_n_ (n ≥ 6) if PTS occurs when copying the opposite strand. As a consequence, an extra A (or U) is added in a fraction of the newly synthetized RNA molecules, so that the gene overprinted in the -1/+2 frame becomes accessible when edited transcripts are translated. Importantly, it has been observed that viral RNAs featuring a one-nucleotide (-nt) deletion at the slippage motifs (A_n_ -> A_n-1_, with n ≥ 6), are also generated in potyviruses, albeit at very low levels (11). Moreover, these one-nt-shorter RNAs might translate the +1 frame into truncated, yet functional proteins, as observed in the case of P3N-ALT from clover yellow vein virus (ClYVV) (13).

With more than 200 assigned representatives sorted in 12 different genera, the *Potyviridae* family is the largest and most socio-economically relevant group of plant-infecting RNA viruses. Potyvirids (members of the family *Potyviridae*) have monopartite (except those in the *Bymovirus* genus, which are bipartite), single stranded and positive sense (+ssRNA) genomes of around 10kb (14–16). It was assumed for a long time that potyvirid genomic RNAs translated only a single, full size, ORF to produce a large polyprotein further processed by viral-encoded proteases into the mature viral factors, usually: P1, HCPro, P3, 6K1, CI, 6K2, NIa (VPg-NIaPro), NIb and CP with some variations mostly concentrated in the N-terminal part of the polyprotein (17). However, since 2008, it has been established that an additional ORF, known as *pipo*, is overprinted in the -1/+2 frame of the P3 coding region. This ORF is preceded by a conserved GA_6_ motif and is expressed as a fusion to the N-terminal half of P3, resulting in the production of P3N-PIPO (18), a protein required for viral movement between cells (19, 20). Few years later, two independent groups discovered that the *pipo* ORF is accessed in potyviruses through PTS at the GA_6_ motif (8, 9). Subsequent investigations have revealed the existence of another overprinted ORF, referred to as *pispo*, located in the -1/+2 frame of the P1 coding sequence in potyviruses infecting sweet potato (21, 22). As for *pipo*, this ORF is accessed by PTS at a conserved GA_6_ motif. Consequently, the edited transcript is translated as a fusion to the N-terminal half of P1, resulting in the production of P1N-PISPO -a protein exhibiting RNA silencing suppression activity (8, 10, 23).

Among well-defined examples of PTS in RNA viruses, the expression of overprinted genes in potyvirids stands out as a unique opportunity to study this phenomenon. Currently, only a limited number of PTS events in viruses belonging to the *Potyvirus* genus within the *Potyviridae* family have been experimentally validated. Based on reported data, PTS in plant RNA viruses exhibits the following characteristics: (i) it is observed in potyviruses and is expected to occur in all members of the *Potyviridae* family due to the conservation of the *pipo* ORF embedded in P3 coding sequences; (ii) it depends on the presence of a GA_n_ (n ≥ 6) motif; and (iii) it more frequently results in a single nucleotide insertion (SNI; A_n_ - > A_n+1_, with n ≥ 6) than a single nucleotide deletion (SND; A_n_ - > A_n-1_, with n ≥ 6). Our study, which encompasses the analysis of PTS in various potyvirids from both the *Ipomovirus* and *Potyvirus* genera, and even in the unrelated potato virus X (PVX, *Potexvirus* genus, *Alphaflexiviridae* family), expands our understanding of this mechanism. Overall, our results highlight the huge flexibility of RNA viruses in exploring the expression of alternative and truncated ORFs by exploiting PTS.

## RESULTS

### An atypical PTS in the ipomovirus CocMoV

Although the GA_6_ slippage motif is present in the central region of the P3 coding sequence of all viruses in the *Potyviridae* family (very few slightly different sequences were recently reported, see below), empirical demonstration of PTS has only been reported in members of the *Potyvirus* genus, such as PPV, ClYVV, turnip mosaic virus (TuMV), sweet potato feathery mosaic virus, bean common mosaic virus, bean common necrotic mosaic virus (8–10, 13, 23). This lack of PTS evidences in viruses from other genera, as well as the presence of an additional GA_6_ motif in its P1 coding sequence (Figure 1A), prompted us to analyse PTS in CocMoV, which belongs to the *Ipomovirus* genus in the family *Potyviridae*. To do that, symptomatic upper non-inoculated leaves of melon plants infected with a natural isolate of CocMoV were harvested at 20 days post-inoculation (dpi) and their RNAs were further used to get amplicons spanning both slippage motifs by RT-PCR (Figure 1A). It is worth noticing here that, unlike in the case of *pipo*, if slippage happens at the GA_6_ motif located in the P1 coding sequence, it will not get access to an overprinted ORF due to stop codons just downstream of the GA_6_ in both alternative frames. Instead, a truncated version of P1, hereafter called P1aN-ALT for its similarity with the truncated P3N-ALT from ClYVV (13), would be translated (Figure 1A).

**Figure 1.**
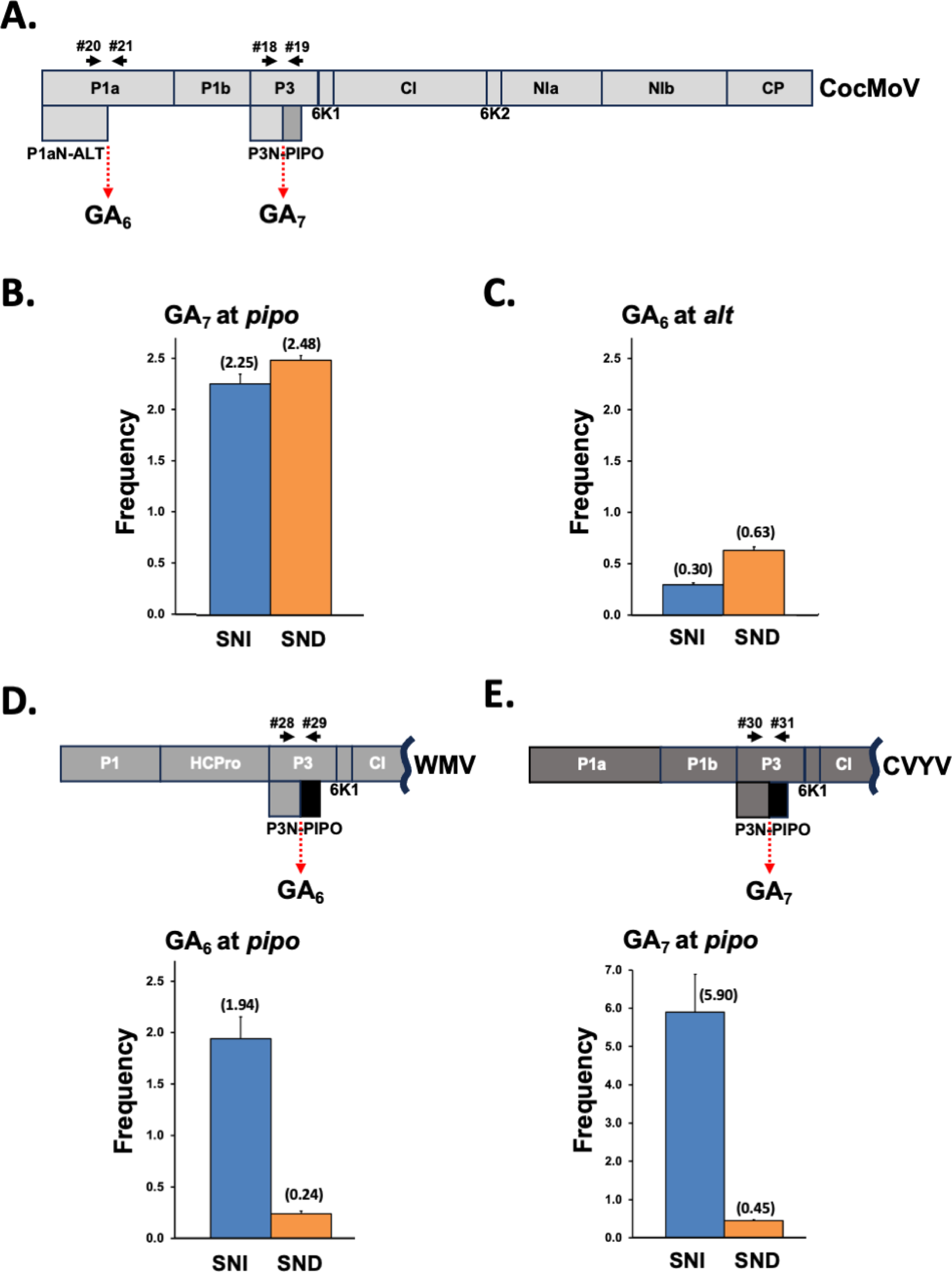
Slippage rates of potyvirids from different genera in melon plants. (A) Schematic representation of the CocMoV coding sequence. (B) Slippage rates at the GA_7_ motif located upstream of the *pipo* ORF from CocMoV. (C) Slippage rates at the GA_6_ motif in *alt* from CocMoV. (D) Schematic representation of a WMV partial coding sequence (upper panel). Slippage rates at the GA_6_ motif located upstream of the *pipo* ORF from WMV (bottom panel). (E) Schematic representation of a CVYV partial coding sequence (upper panel). Slippage rates at the GA_7_ motif located upstream of the *pipo* ORF from CVYV (bottom panel). Bars represent the average frequencies (indicated between brackets) for single nucleotide insertions (SNI) and single nucleotide deletions (SND) at the indicated slippage motifs. Error bars represent standard deviation (n = 3). Primers used to get amplicons are represented with black arrows in the virus map.

The analysis by NGS of amplicons covering the GA_6_ motif right upstream of the *pipo* ORF in CocMoV revealed that insertional slippage, detected by the presence of an additional A at the slippage motif (GA_6_ -> GA_7_), is observed in this ipomovirus at 2.30% of the sequenced molecules (Figure 1B). Surprisingly, this analysis also revealed that a similar number of molecules, 2.48%, suffered deletional slippage, observed as the loss of one A at this position (GA_6_ -> GA_5_) (Figure 1B). Although SND at the slippage motif has been already reported in potyviruses, they were clearly underrepresented in all cases when compared with SNI, and even considered as the result of possible artefacts produced during the preparation of amplicons (9).

The analysis of PTS at the additional GA_6_ motif in CocMoV, located in the P1 coding sequence (Figure 1A), allowed us to test whether slippage also takes place at a different position of the viral genome and, if so, whether SND and SNI are represented by similar frequencies as found at the motif that precedes the *pipo* ORF. At this particular site, we also found slippage, but at a lower overall frequency: 0.93% (Figure 1C). Remarkably, species with SNI represented just 0.30% of the sequenced molecules, whereas those with SND were 0.63% (Figure 1C).

To test whether such a high frequency for SND in CocMoV was a distinctive feature of this particular virus, as well as to discard potential artefacts, we also inoculated melon plants with other two potyvirids, WMV (genus *Potyvirus*) and CVYV (genus *Ipomovirus*), in order to estimate PTS. Upper non-inoculated leaves of infected plants showing clear disease symptoms were harvested at 24 and 10 dpi for WMV and CVYV, respectively, and their RNAs were used to get amplicons spanning their corresponding slippage motifs upstream of the *pipo* ORF by RT-PCR, which were further subjected to NGS. As expected, PTS events were detected in both viruses at this position. In the case of WMV, the rate of SNI was 1.94%, which was eight times higher than the 0.24% corresponding to SND (Figure 1D). A similar result was observed in CVYV, which belongs to the same genus as CocMoV. Here, the overall slippage was higher: 5.90% for SNI and 0.45% for SND, resulting in a SNI 13 times higher than SND (Figure 1E). Therefore, we conclude that PTS in CocMoV differs from other potyvirids in having a high frequency of SND at slippage sites, which is equivalent, or even higher, than that of SNI.

### The atypical PTS in CocMoV is not the consequence of nucleotides flanking slippage motifs

In a previous study, Olspert and collaborators demonstrated that the overall slippage frequency in TuMV is affected by nucleotides flanking the minimal GA_6_ motif (11). If these nucleotides were responsible for the atypical PTS in CocMoV, then RNA segments spanning the slippage sites of CocMoV would induce CocMoV-like slippage in a heterologous potyvirid. To test this hypothesis, we chose PPV given that PTS in this virus has been already studied and showed the typical SNI >> SND (8). We first tested whether PPV supports a genuine second slippage motif by introducing the most slippery, synthetic, G/C rich 21-nt sequence based on a GA_6_ central motif developed by Olspert and collaborators (named PISPO 5’&3’str in (11) and HF_slip in this study). This segment was introduced upstream of an out-of-frame eGFP coding sequence to generate an infectious clone that produces PPV-HF_slip_eGFP, which would express eGFP if a +1 frameshift took place (Figure 2A). In addition, we constructed an equivalent PPV derivative in which the “AAAAAA” in HF_slip was mutated to “AGAAGA” in order to generate an infectious clone that produces PPV-HF_slip_mut_eGFP, where no slippage would be expected. When inoculated in *N. benthamiana,* the control PPV-eGFP expressed high levels of eGFP in upper non-inoculated leaves at 9 dpi, as observed by both fluorescent microscopy and GFP immunodetection, whereas PPV-HF_slip_eGFP expressed, although at low levels, this reporter, suggesting that PTS was taking place (Figure 2B). In contrast, PPV-HF_slip_mut_eGFP did not display fluorescent nor chemiluminescent signals derived from eGFP (Figure 2B), supporting the idea that PTS was abolished when the A_6_ stretch is mutated. NGS of RT-PCR products spanning the motifs upstream of the *egfp* ORF confirmed these observations: slippage was not observed at HF_mut_slip (not shown), but clearly detected at the HF_slip motif: 3.88% for SNI and 0.48% for SND (Figure 2C). As expected from previous results (8), the same trend with SNI >> SND was observed at the natural slippage motif located upstream of the *pipo* ORF: 2.59% and 0.13%, respectively (Figure 2C). With these data, we conclude that PPV is able to support the expression of two independent out-of-frame products via genuine PTS. NGS analyses of amplicons made by PCR directly with the plasmid as template (negative control) showed barely detectable levels of SNI, and very low levels of SND, which were comparable to those observed in RT-PCR products, especially in the HF_slip motif (Figure 2C). Very low but still detectable rates of SND in PCR products made directly from plasmids have been previously reported, suggesting that SND events occur at a low frequency during amplicon preparation and sequencing (11).

**Figure 2.**
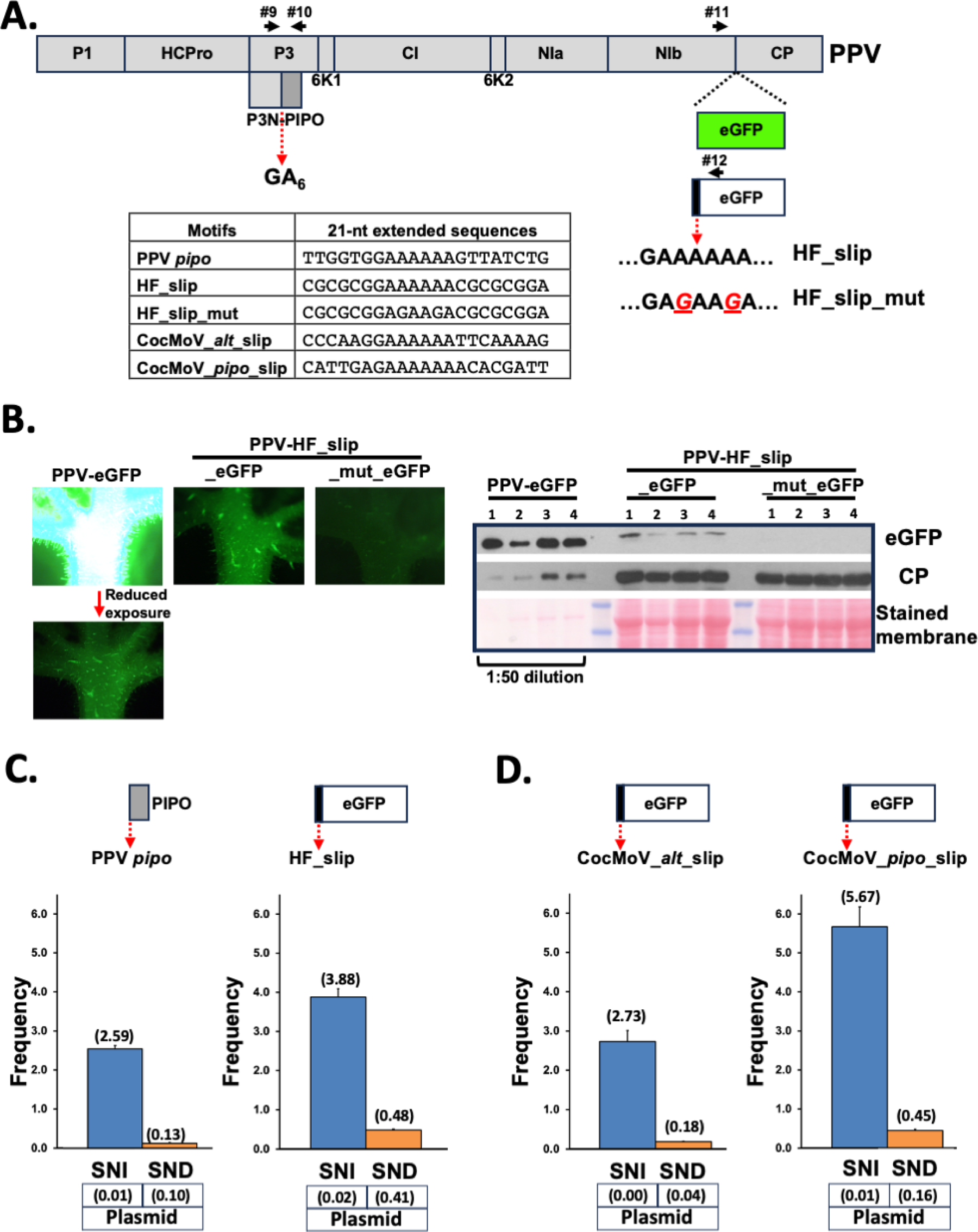
Expression of an overprinted reporter gene via PTS at diverse motifs in the context of PPV infection. (A) Schematic representation of the PPV coding sequence. The insertion of eGFP and the out-of-frame eGFP coding sequences in the virus genome are shown. The 21-nt slippage motifs introduced upstream of the out-of-frame eGFP coding sequence are indicated in the table. Primers used to get amplicons are represented with black arrows. (B) Representative pictures taken under UV radiation at 9 days post-inoculation of upper non-inoculated *N. benthamiana* leaves infected with the indicated viruses are shown at the right. The leaf infected with PPV-eGFP had to be exposed 100 times less to UV radiation to display equivalent fluorescent signal than the leaf infected with PPV-HF_slip_eGFP. The lefts panel shows the detection of eGFP and PPV CP by immunoblot analysis in protein samples from upper non-inoculated leaves of four *N. benthamiana* plants infected with the indicated viruses. Blot stained with Ponceau red showing the large subunit of the ribulose-1,5-bisphosphate carboxylase-oxygenase is included as a loading control. Protein extract from leaves infected with PPV-eGFP had to be diluted 50 times to display comparable eGFP-derived chemiluminescent signal. (C) Slippage rates at the GA_6_ motif located upstream of the *pipo* ORF from PPV (left panel). Slippage rates at the HF_slip motif located upstream of the *egfp* ORF inserted in the PPV RNA (right panel). (D) Slippage rates at the CocMoV_*alt*_slip motif located upstream of the *egfp* ORF inserted in the PPV RNA (left panel). Slippage rates at the CocMoV_*pipo*_slip motif located upstream of the *egfp* ORF inserted in the PPV RNA (right panel). Bars represent the average frequencies (indicated between brackets) for single nucleotide insertions (SNI) and single nucleotide deletions (SND) at the indicated slippage motifs. Error bars represent standard deviation (n = 3). The slippage rates obtained at each of those motifs by using an amplicon directly produced by PCR from the cloned viral cDNA are indicated (plasmid).

Finally, we replaced the HF-slip motif that precedes the *egfp* ORF in PPV-HF_slip_eGFP by two independent segments of 21 nucleotides corresponding to both CocMoV slippage sites. Upper non-inoculated leaves of infected *N. benthamiana* plants were harvested at 9 dpi and their RNAs were used to get amplicons spanning the motifs preceding the *egfp* ORFs by RT-PCR. Analyses of SNI and SND rates indicated that both slippage motifs from CocMoV promoted, in the context of a PPV infection, much more SNI than SND (Figure 2D), ruling out the possibility that the atypical PTS in CocMoV is due to nucleotide sequences flanking the slippage motifs. NGS of PCR products made from plasmids (negative controls) showed, once again, just very low levels of SND (Figure 2D). As in the context of CocMoV infection (Figure 1B and C), the overall slippage frequency at the motif upstream the *pipo* ORF was higher than that at *alt*. Remarkably, it was even higher than that at the HF_slip motif, which was indeed selected for this study as it promoted the highest slippage frequency out of many other tested motifs (11). These differences can be explained by the direct correlation between the slippage frequency and the length of homopolymeric runs of As (A_7_ for CocMoV_pipo_slip, and A_6_ for CocMoV_alt_slip and HF_slip), as observed for a modified slippage motif that carries either six, seven or eight consecutive As (11).

### The atypical PTS in CocMoV can be attributed to particular features of CocMoV

Based on results described above, we hypothesized that intrinsic features in CocMoV unrelated to its genomic RNA sequence define its peculiar PTS. To test this hypothesis, we carried out the reciprocal experiment to the one showed in Figure 2 by modifying the CocMoV genome in order to introduce additional slippage motifs that could induce much more SNI than SND. First, we built an infectious cDNA clone for CocMoV suitable to conveniently manipulate the viral genome (see Materials and Methods for more details). Zucchini and melon plants (n = 6 per plant species) were inoculated with the generated infectious clone by bombardment, and the infection process was monitored over the time. All plants started to display the expected symptoms of CocMoV infection in upper non-inoculated leaves at 10 dpi, like the natural isolate, and the presence of CocMoV in these tissues was confirmed by RT-PCR (data not shown). The CocMoV infectious clone was further manipulated to independently introduce, in the P1 coding sequence, and in frame with the main virus ORF, 21-nt fragments comprising the HF_slip motif and the slippage sequence upstream of the PPV *pipo* ORF (Figure 3A). Symptomatic upper non-inoculated leaves of melon plants infected with CocMoV derivatives were harvested at 20 dpi and their RNAs were used to get amplicons spanning diverse slippage motifs by RT-PCR, which were further subjected to NGS. The analysis of natural slippage motifs in the CocMoV derivative that carries HF_slip at the 5’end of its genome confirmed our results with the wild type virus (Figure 1B and C), as rates for SNI and SND were comparable at the GA_7_ motif that precedes the *pipo* ORF (Figure 3B), whereas the SND rate was higher than that of SNI at the GA_6_ motif located at the 3’end of *p1an-alt.* Importantly, slippage rates produced at the HF_slip introduced in the 5’ end of CocMoV genome, contrary to what we found in the context of PPV infection, were similar to those at the CocMoV GA_6_ motif in the 3’end of *p1an-alt*, displaying SND >> SNI (Figure 3C). A more remarkable difference between SND and SNI was observed with a CocMoV derivative that carries, at the 5’ end of its genome, the slippage motif located upstream of the PPV *pipo* ORF (Figure 3D). We also analyzed by NGS equivalent amplicons obtained directly by PCR reactions using plasmids as template (negative controls). Similar to what we observed in amplicons obtained directly from PPV-based plasmids (Figure 2C and D), small rates of SND were detected (Figure 3C and D), but these cannot explain, in any way, the high levels of SND observed in amplicons obtained from infected plants by RT-PCR. Having in mind that CVYV, a close relative of CocMoV that shares a similar host range and belongs to the same *Potyviridae* genus (*Ipomovirus*), displays the typical SNI >> SND when infecting melon (Figure 1E), the most likely explanation for the peculiar PTS found in CocMoV is that its viral polymerase is more prone to delete one nucleotide when replicating homopolymeric runs of As. As far as we know, this viral feature has not been described before.

**Figure 3.**
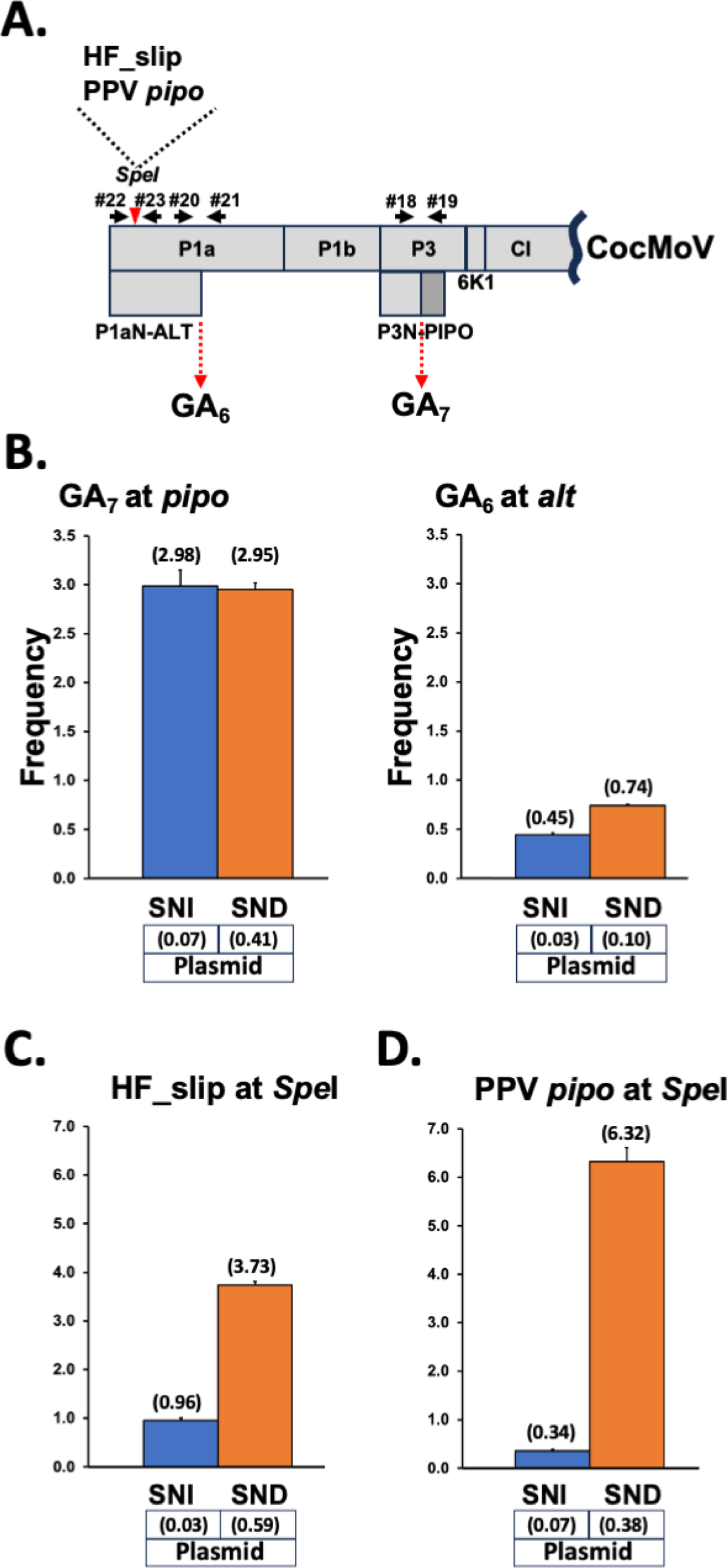
PTS promoted by artificial slippage motifs in the context of CocMoV infection. (A) Schematic representation of a CocMoV partial coding sequence. The unique SpeI restriction site, in which the indicated 21-nt slippage motifs were introduced, is indicated with a red triangle. Primers used to get amplicons are represented with black arrows. (B) Slippage rates at the GA_7_ motif located upstream of the *pipo* ORF from CocMoV (left panel). Slippage rates at the GA_6_ motif located upstream of *alt* in CocMoV (right panel). (C) Slippage rates at the HF_slip motif inserted in the unique SpeI site of CocMoV coding sequence (D) Slippage rates at the PPV *pipo* motif inserted in the unique SpeI restriction site of CocMoV coding sequence. Bars represent the average frequencies (indicated between brackets) for single nucleotide insertions (SNI) and single nucleotide deletions (SND) at the indicated slippage motifs. Error bars represent standard deviation (n = 3). The slippage rates obtained at each of those motifs by using an amplicon directly produced by PCR from the cloned viral cDNA are indicated (plasmid).

### Wide distribution of diverse slippage motifs in the family *Potyviridae*

It has been already shown in a potyvirus, ClYVV, that a truncated P3 protein, produced from transcripts carrying 1-nt deletion in the *pipo* A_6_ stretch, is required for viral movement (13). This result, along with our unexpected findings with CocMoV, support the idea that not only overlapping ORFs acceded by +1 frameshift have to be considered as potential functional products, but also overlapping ORFs acceded by -1 frameshift, as well as truncated versions of proteins harbouring coding sequences with homopolymeric runs of As. Additionally, in view of previous results from Olspert and collaborators showing that an artificial slippage motif with U_6_ (instead of A_6_) also induces PTS in TuMV (11), it seems likely that the number of functional factors produced by viruses of the *Potyviridae* family is underestimated.

Based on the above premises, we searched for A_n_/U_n_ (n ≥ 6) stretches in the coding sequence of fully sequenced individuals in the *Potyviridae* family (n=189 species in the latest ICTV release, 13^th^ of September 2023, https://ictv.global/vmr). Sequences features are summarized in Supplementary Table S1. A first remarkable observation is that only two potyvirids, brugmansia suaveolens mottle virus (BsMoV, GenBank accession AB551370) and johnson grass mosaic virus (JGMV, GenBank accession Z26920), do not have any A_n_/U_n_ (n ≥ 6) string in their genomes. However, after a careful inspection, we found that they do harbour a well-defined overlapping *pipo* ORF embedded in their P3 coding sequence (data not shown), which is preceded in both cases by GA_5_U instead of the common GA_6_ motif. The analysis of available genome sequences of more recently sequenced JGMV isolates (GenBank accessions MZ405658, KT833782 and KT289893) show, instead, that a GA_6_ motif is located upstream of their *pipo* ORF. On the other hand, and thanks to Alice Kazuko Inoue-Nagata and her team, we were able to revisit the original Sanger sequencing data of the only reported isolate of BsMoV (39). We found that two out of the three sequenced clones harbouring the RT-PCR product from this virus have, indeed, GA_5_U upstream of the *pipo* ORF, but the remaining one (deposited recently with GenBank accession OR497357) has a GA_6_ slippage motif at that position. Given that the genome sequence of BsMoV, as well as that of the first reported JGMV, was assembled from few cloned RT-PCR products (39, 40), we envisage two putative scenarios to explain the lack of slippage motifs preceding the *pipo* ORF in the reported full-length sequences of these two viruses: (i) the presence of GA_5_U is simply an artefactual mutation introduced during RT-PCR, or (ii) plants might be infected with a mix of viruses in some cases, including variants without the conserved GA_6_ slippage motif right upstream of the *pipo* ORF. If the second option were true, then further studies would be required to understand the biological meaning of it.

Another observation is that slippage motifs preceding the *pipo* ORF in 9 out of 189 fully-sequenced potyvirids are either CA_6_ (six cases) or UA6 (three cases) instead of GA_6_ (Supplementary Table S1). Moreover, two recently reported potyvirids, tentatively named stylo mosaic-associated virus 1 (StyMaV1) and StyMaV2, not yet included in the last list released by ICTV, also have CA_6_ and UA_6_ slippage motifs, respectively, immediately upstream of the *pipo* ORF (41). Overall, these findings indicate that G at the first position of functional slippage motifs is dispensable for PTS to happen in nature.

The Supplementary Table S1 also shows that, although the majority of potyvirids (130 out of 189) only have the slippage motif required to get access to the *pipo* ORF (nucleotide positions highlighted in bold and underlined), some others have additional A_n_/U_n_ (n ≥ 6) signatures in their genome with the potential to produce truncated forms of certain proteins in most cases (nucleotide positions are indicated), or even to get access to short alternative ORFs with more than 90 nucleotides (nucleotide positions are indicated and highlighted in bold). Additional studies will be required on a case-by-case basis to define the relevance of such additional slippage motifs over viral infections.

### A naturally-occurring U_8_ motif promotes access to the *pipo* ORF

Surprisingly, the only available full-length genome of WMVBV (isolate Su03-07) carries no A_n_ (n ≥ 6) motif at the P3 coding sequence. Instead, it has a U_8_ stretch located 16 nucleotides upstream of a well-defined overlapping *pipo* ORF (Figure 4A and Supplementary Table S1). We also determined the nucleotide sequence of a short fragment of the P3 coding region from Su12-21, another isolate of WMVBV for which the complete genome sequence is not available (25). A blast analysis of the WMVBV Su12-21 nucleotide sequence only retrieved a fragment of WMVBV Su03-07, and indicated that the U_8_ motif is conserved in both isolates, with an overall identity of 89% for the whole sequenced fragment (Supplementary Figure S2). Based on these findings, we hypothesized that PTS might take place at the U_8_ stretch in the WMVBV genome during viral replication to get access to the *pipo* ORF. To test this idea, upper non-inoculated leaves of melon plants infected with these two WMVBV isolates were harvested at 30 dpi, and amplicons spanning the slippage motif were obtained from total RNA samples by RT-PCR. NGS analysis of these amplicons revealed that, as anticipated, PTS events are clearly observed for both isolates at this position (Figure 4B). Remarkably, the rate of SNI and SND were similar in each isolate: 4.72% and 3.80% for Su03-07, and 6.22% and 4.27% for Su12-21. Differences in SNI and SND rates when comparing Su03-07 versus Su12-21 resulted significative (*p* value < 0.01, Figure 4B), and it might be due to variations in the genome sequence around the slippage motif. In fact, the Su12-21 isolate, which displayed higher slippage frequencies, has two Gs just upstream of the U_8_ motif. In turn, Su03-07 has two As in these particular positions (Supplementary Figure 2), which is in perfect agreement with previous findings indicating that increased flanking sequence GC content upstream and/or downstream of the homopolymeric run correlates with higher PTS rates (11).

**Figure 4.**
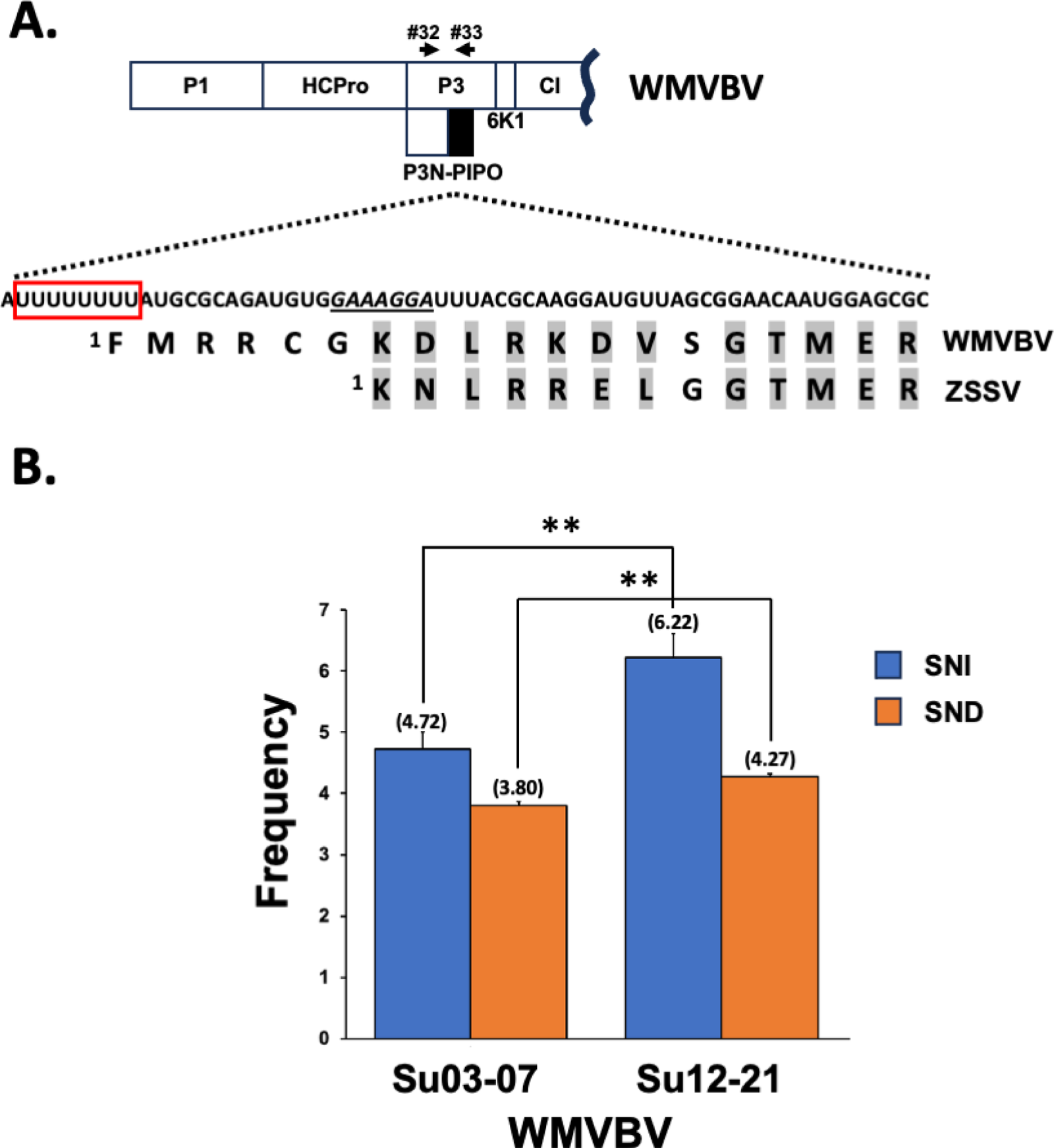
The unusual slippage motif located upstream of the *pipo* ORF in WMVBV. (A) Schematic representation of a WMVBV partial coding sequence. Nucleotide and amino acid sequences corresponding to the N-terminal part of PIPO in WMVBV are depicted. A sequence of nucleotides reminiscent to a GA_6_ motif is underlined. A U_8_ motif is highlighted with a red box. A partial alignment of PIPO from WMVBV and ZSSV, in which equivalent/related amino acids are shadowed in grey, is shown. Primers used to get amplicons are represented with black arrows. (B) Slippage rates at the U_8_ motif located upstream of the *pipo* ORF from the indicated WMVBV isolates. Bars represent the average frequencies (indicated between brackets) for single nucleotide insertions and single nucleotide deletions. Error bars represent standard deviation (n = 3). Statistical differences were tested with the post-hoc Tukey HDS test (** *p* value < 0.01).

So far, all sequenced potyvirids, with the only exception of WMVBV, have A_6_ motifs preceding the *pipo* ORF (Supplementary Table S1). In this scenario, the simplest evolutionary pathway that explains this observation is that a common ancestor of all potyvirds had A_6_ upstream of the *pipo* ORF, and that the U_8_ motif evolved during WMVBV speciation. Indeed, a careful inspection of WMVBV genome at the slippage region, and of the PIPO amino acid sequence in zucchini shoestring virus (ZSSV), a virus closely related to WMVBV (15), provides some clues supporting this idea. As observed in Figure 4A, the first amino acid in ZSSV PIPO is a K, which aligns with the seventh amino acid of WMVBV PIPO. As the WMVBV coding sequence at that particular position vaguely resembles the conserved GA_6_ motif (Figure 4A), we propose that a WMVBV ancestor evolved a U-rich stretch few nucleotides upstream of the *pipo* ORF, promoting higher slippage rates than the original GA_6_ motif with a potential improvement in cell-to-cell movement. In fact, from all the natural PTS motifs preceding the *pipo* ORF analysed in this study, the U_8_ from WMVBV is the one displaying the highest SNI + SND rates. Another non-mutually exclusive possibility is that the presence of six extra amino acids at the N-terminus of WMVBV PIPO (FMRRCG, Figure 4A) improves the performance of P3N-PIPO.

### Many slippage motifs in a single potyvirid

As shown in Supplementary Table S1, one interesting potyvirid is ANSSV (*Arapavirus* genus), as its only known genome sequence has the highest number of A_n_/U_n_ (n ≥ 6) motifs. Aiming to know whether PTS takes place in all of these motifs, we carried out a transcriptomic analysis of leaves from an areca palm infected with ANSSV. With a mean coverage of 34000X per nucleotide position, we were able to estimate both SNI and SND frequencies with high confidence. For this analysis we considered that background slippage rates for SNI and SND are 0.28% (at position 2028) and 0.63% (at position 7088), respectively, due to the slippage events observed in sequences different from A_n_/U_n_ (n ≥ 6) (positions 2028 and 7088) (Figure 5). Therefore, whereas no peak above the background was observed for SND, three positions presented a rate of SNI way above the background (Figure 5). A careful inspection of those positions indicated that they correspond to three A_n_ (n ≥ 6) motifs including, as expected, the one immediately upstream of the *pipo* ORF (position 2622, Figure 5).

**Figure 5.**
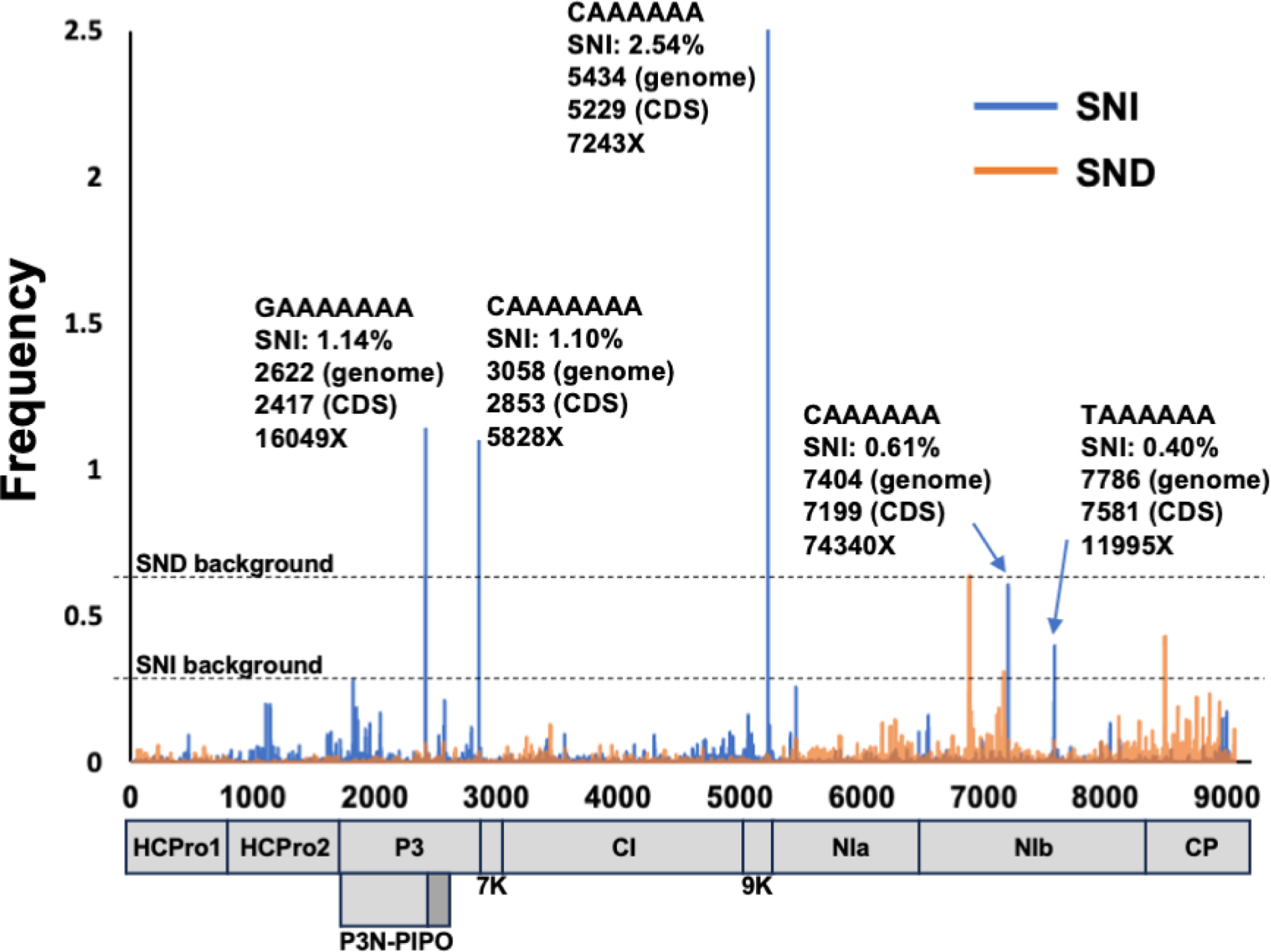
Many slippage motifs in the genome of ANSSV. Schematic representation of the ANSSV coding sequence. Rates of single nucleotide insertions (SNI) and single nucleotide deletions (SND) in the whole ANSSV genome are shown. Slippage motifs associated to each peak along with its SNI rate, position in the viral genome and viral coding sequence, as well as the sequencing coverage at particular position, are indicated.

Even more interesting are the following findings:

i. the A_7_ motif upstream of the *pipo* ORF (position 2622) is not the one displaying the highest PTS frequency. Indeed, an A_6_ motif at position 5434 promoted higher levels of SNI (2.54% vs 1.14%, Figure 3). This particular motif, along with another relevant A_7_ run at position 3058 (SNI = 1.10%, Figure 5), gives no access to overlapping ORFs, so that they might be playing another role. It is relevant to mention that both motifs are located around the boundaries of two cistrons (P3-7K for the motif at position 3058, and 9K-NIa for that at position 5434), and one can speculate that they play a role in the regulation of polyprotein truncation at certain places via the introduction of premature stop codons;
ii. the only presence of a A_n_/U_n_ (n ≥ 6) motif in a potyvirid is not enough for significant PTS to happen. ANSSV has seven of these motifs in its genome, those mentioned in point (i), three additional A_6_ runs and one U_6_ motif. From the last four, the A_6_ motifs at position 7404 and 7786 displayed a PTS frequency just above the background (SNI of 0.61% and 0.40, respectively, Figure 5). However, runs of A_6_ at position 5665 and U_6_ at position 5619 displayed a marginal SNI of 0.25% and 0.10%, respectively.

Diversity in promoting PTS at different rates when comparing the seven motifs might be due to differences in the flanking nucleotide sequences, as previously shown with artificially-introduced slippage motifs in TuMV (11). All in all, data presented so far highlight the sequence flexibility of natural slippage motifs (long homopolymeric runs of As or Us), and reinforce the idea that PTS might take place at different positions, all along potyvirid genomes, independently of the presence/absence of downstream overlapping ORFs.

### Factors unrelated to the cognate virus shape PTS rate

#### Effect of mixed infections on PTS rate

To test whether the presence of another virus alters SNI and/or SND rates in CocMoV, we performed mixed infections of CocMoV + WMV and CocMoV + CYSDV. We harvested systemically infected leaves at 20 dpi and prepared RNA samples in order to get amplicons spanning CocMoV slippage motifs by RT-PCR, which were then subjected to NGS. We found that slippage rates at the GA_6_ motif that precedes *p1an-alt* in CocMoV were not affected by the presence of WMV (Supplementary Figure S1). On contrary, slippage at the GA_7_ motif that precedes the *pipo* ORF was altered in the mixed infection, as the presence of WMV caused a significative reduction in the SNI rate (2.25% versus 1.86%) (Supplementary Figure S1). Different results were observed in plants infected with CocMoV + CYSDV, as in this particular case both SNI and SND were slightly higher at the GA_7_ motif that precedes the *pipo* ORF (2.86% and 2.63% versus 2.25% and 2.48%, respectively) (Supplementary Figure S1). Comparable to the case of CocMoV + WMV, PTS rates at the GA_6_ motif that precedes the *p1an-alt* ORF were not significatively affected in CocMoV + CYSDV infected plants (Supplementary Figure S1). All in all, these results are in line with our previous observation in a different pathosystem (23), so confirming that coinfection with other viruses can affect PTS rates. Even though additional experiments are required to understand the meaning of these observations, as well as mechanisms underpinning these effects, we speculate that complementation and competition are taking place.

#### Host effect on PTS

To test whether SNI and SND rates for a given virus are somehow influenced by the host identity, we used SPV2 (*Potyvirus* genus) as model given that:

i. SPV2 belongs to the subgroup of sweet potato-infecting potyviruses, which encode two ORFs overlapping the main viral ORF: *pispo* in the P1 coding sequence and *pipo* in that of P3 (Figure 6A), both acceded by PTS as demonstrated in the closely-related sweet potato feathery mottle virus (SPFMV) (8, 10),
ii. SPV2 is able to infected diverse hosts besides ipomeas (42), such as the model plant

**Figure 6.**
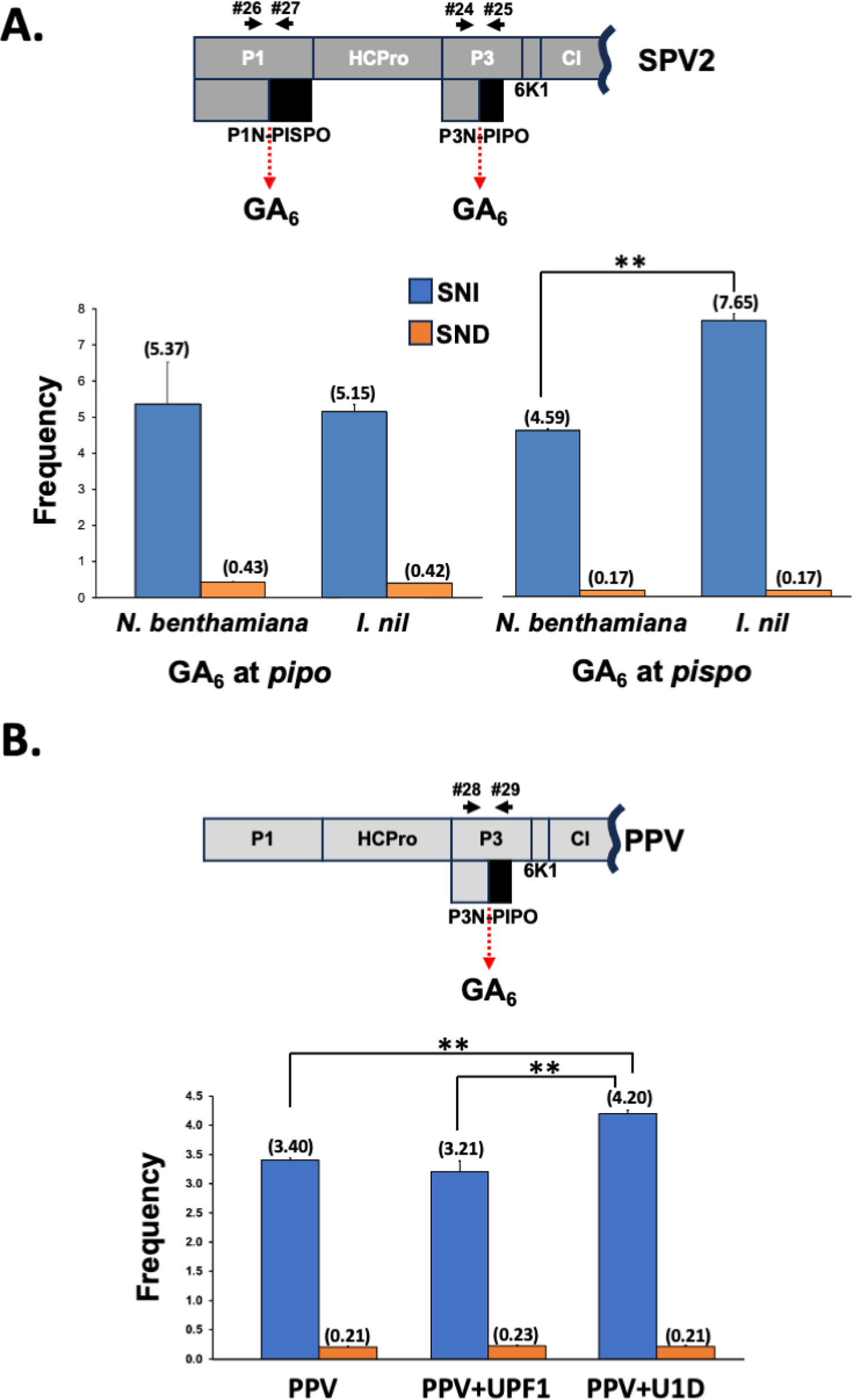
Host influences on PTS rates. (A) Schematic representation of a SPV2 partial coding sequence, including slippage motifs and primers used to get amplicons (black arrows). Slippage rates at the GA_6_ motifs upstream both *pipo* and *pispo* ORFs from SPV2 are shown. (B) Schematic representation of a PPV partial coding sequence, including the slippage motif and primers used to get amplicons (black arrows). Slippage rates at the GA_6_ motif upstream of the *pipo* ORF from PPV in PPV-infected plants, supplemented in *trans* with the indicated proteins by agroinfiltration, are shown. Bars represent the average frequencies (indicated between brackets) for single nucleotide insertions (SNI) and single nucleotide deletions (SND) at the slippage motifs. Error bars represent standard deviation (n = 3). Statistical differences were tested with the post-hoc Tukey HDS test (** *p* value < 0.01).

#### N. benthamiana

Symptomatic upper non-inoculated leaves of *Ipomea nil* (natural host) and *N. benthamiana* (laboratory host) plants infected with SPV2 were harvested at 14 dpi, and the corresponding RNA samples were used to get amplicons spanning both slippage motifs by RT-PCR (Figure 6A). NGS analysis of amplicons showed that, as expected, PTS took place during viral replication at both motifs in both hosts. Like for most viruses studied here and in previous works, SNI rates were much higher than those of SND (Figure 6B and C). Interestingly, although PTS rates at the motif located immediately upstream of the *pipo* ORF were similar in both hosts (Figure 6B), those at the motif that precedes the *pispo* ORF were significatively different, particularly in the case of SNI (*p* value < 0.01). SNI was more frequent in *I. nil* than in *N. benthamiana* (7.65% versus 4.59%, Figure 6C), supporting the idea that P1N-PISPO, the protein expressed from RNA molecules carrying an additional A, accumulates at higher levels in the natural host *I. nil*. Assuming that SPV2 P1N-PISPO interferes with host RNA silencing like its counterpart from SPFMV (10, 23), and given that the laboratory strain of *N. benthamiana* is a natural mutant of RDR1, a key protein for host-based defence mediated by RNA silencing (43, 44), one can speculate that the pressure to produce P1N-PISPO is lower in this host; in this scenario, slippage at the motif that precedes the *pispo* ORF would not be as necessary in *N. benthamiana* as in other silencing proficient plants.

#### Host-dependent nonsense-mediated RNA decay affects PTS rate

A direct consequence of PTS in potyvirids is the production of viral RNAs with an exceptionally large 3’ untranslatable region (UTR). For instance, RNA species coding for P1-HCPro-P3NPIPO harbour 3’ UTRs of around 7000 nucleotides. In all eukaryotes, coding RNAs with large 3’ UTR are commonly targeted by the nonsense-mediated mRNA decay (NMD), which is a surveillance pathway that safeguards the quality of the transcriptome by eliminating mRNAs with premature stop codons (45). In order to test whether PTS products are recognized and eliminated by the NMD pathway, we infected *N. benthamiana* plants with PPV and further blocked NMD in infected, upper non-inoculated leaves. Such a blockage was achieved by inhibiting UPF1, a key factor of the NMD mechanism, via the expression of the dominant negative mutant U1D (30). As controls, we used untreated infected plants, as well as plants transiently expressing the wild type UPF1. Three days after the treatment, total RNAs were prepared from systemically infected leaves, which were further used as template to get amplicons spanning the PPV slippage motif that precedes the *pipo* ORF by RT-PCR (Figure 2A). NGS analysis of these amplicons showed a significant increase in the SNI rate when the NMD was inactivated (Figure 6D). On the other hand, the analysis also indicated that the very low SND rate does not increase when expressing U1D (Figure 6D). This result might be justified by the previously proposed idea that such a low SND rates are mostly artifacts during library preparation and NGS (11).

#### Programmed transcriptional slippage outside the *Potyviridae* family

Finally, in order to check whether PTS takes place in RNA viruses unrelated to the *Potyviridae* family, we used the well-characterized potato virus X (PVX, *Potexvirus* genus, *Alphaflexiviridae* family) as model. We manipulated a previously described PVX vector (32) to generate an infectious clone that produces PVX-HF_slip-eGFP, a virus that expresses, from a duplicated subgenomic promoter, an out-of-frame eGFP coding sequence. In this case, the reporter is preceded by an AUG (to initiate translation) followed by the HF_slip motif, the eGFP coding sequence and a stop codon (Figure 7A). Hence, as for PPV-HF_slip-eGFP, the eGFP would be produced during infection only if +1/-2 frameshift took place. In addition, we constructed an equivalent PVX derivative in which the A_6_ motif in HF_slip was replaced by AGAAGA to generate an infectious clone that produces PVX-HF_slip_mut_eGFP. These two PVX derivatives, along with PVX-eGFP, in which the eGFP coding sequence is in frame with the initiation codon (Figure 7A), were inoculated in *N. benthamiana* plants and systemically infected leaves were harvested at 6 dpi. As expected, PVX-eGFP expressed high levels of eGFP in upper non-inoculated leaves, as observed by both fluorescent microscopy and GFP immunodetection, whereas PVX-HF_slip_eGFP expressed lower levels of this reporter that was clearly detected by both methods (Figure 7B). In contrast, PVX-HF_slip_mut_eGFP did not display fluorescent nor inmunodetection signals derived from eGFP (Figure 7B).

**Figure 7.**
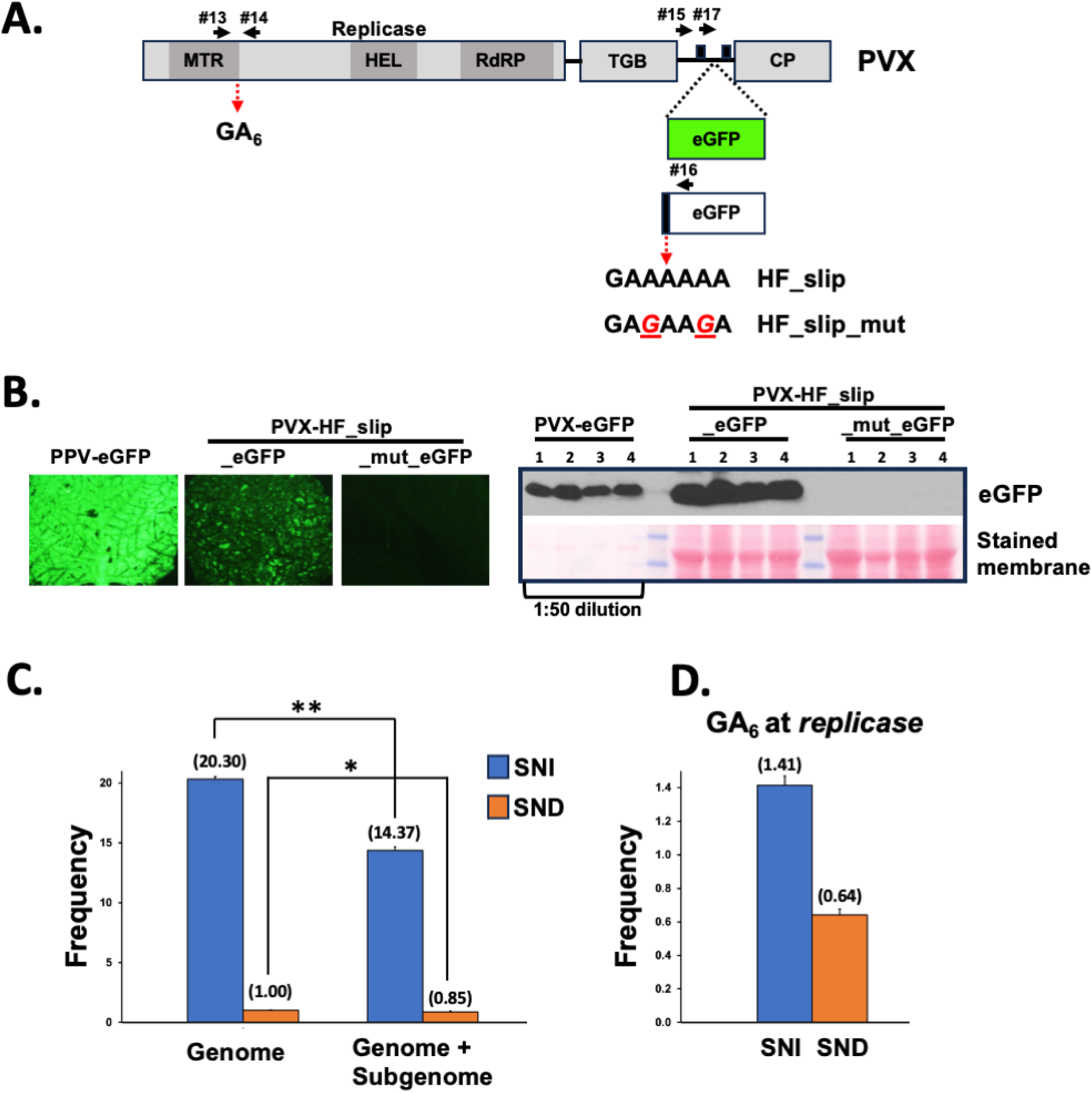
PTS outside of the *Potyviridae* family, the case of PVX. (A) Schematic representation of the PVX coding sequence. The insertion of eGFP and the out-of-frame eGFP coding sequences in the virus genome is shown. The slippage motifs introduced upstream of the out-of-frame eGFP coding sequence are also indicated. The two copies of the subgenomic promoter are depicted as black boxes. Functional domains presented in the replicase protein are indicated (MTR: methyltransferase; HEL: helicase; RdRP: RNA dependent RNA polymerase). Primers used to get amplicons are represented with black arrows. (B) Representative pictures taken under UV radiation at 9 days post-inoculation of upper non-inoculated *N. benthamiana* leaves infected with the indicated viruses are shown at the right. The left panel shows the detection of eGFP by immunoblot analysis in protein samples from upper non-inoculated leaves of four *N. benthamiana* plants infected with the indicated viruses. Blot stained with Ponceau red showing the large subunit of the ribulose-1,5-bisphosphate carboxylase-oxygenase is included as a loading control. Protein extract from leaves infected with PVX-eGFP had to be diluted 50 times to display comparable eGFP-derived chemiluminescent signal. (C) Slippage rates at the HF_slip motif located upstream of the *egfp* ORF inserted in the PVX RNA. (D) Slippage rates at the GA_6_ motif located downstream of the MTR motif from the PVX replicase coding sequence. Bars represent the average frequencies (indicated between brackets) for single nucleotide insertions (SNI) and single nucleotide deletions (SND) at the indicated slippage motifs. Error bars represent standard deviation (n = 3). Statistical differences were tested with the post-hoc Tukey HDS test (* *p* value < 0.05; ** *p* value < 0.01).

The NGS analysis of RT-PCR amplicons spanning the introduced slippage motifs (primer pair #15/#16, Figure 7A) showed background levels of SND and SNI in HF_slip_mut RNAs, confirming that PTS was not taking place in this virus (data not shown). In turn, slippage rates at the HF_slip motif in PVX was even higher than that in PPV: 20.32% for SNI and 1% for SND in the PVX background, versus 3.88% for SNI and 0.48% for SND in the PPV background (Figure 7C and 2C). Such a high PTS frequency in PVX might be due to the particular strategy used to express the overlapping ORF. In the case of PVX, genome variants with +/-A at the slippage motif would retain the propagation potential and are maintained in the viral population as the ORF for the CP cistron, which is located downstream of HF_slip_eGFP, is not disrupted by the upstream editing since the CP transcript is expressed from its own subgenomic promoter (Figure 7A). In contrast, PPV variants with +/-A at the slippage motif would tend to be eliminated from the viral population as the ORF for the CP cistron, which is also located downstream of HF_slip_eGFP, is no longer acceded when PTS takes place (Figure 2A).

To gain insight into the production of overlapping ORFs in PVX, we also carried out NGS analysis of RT-PCR amplicons spanning the introduced slippage motif, but produced with another forward primer (same RNA samples as above, primer pair #17/#16, Figure 7A). Since this primer anneals downstream of the duplicated subgenomic promoter, template for this amplification includes the PVX genome (as above), as well as the subgemonic RNA. As expected, PTS was not detected at significant levels in the HF_slip_mut motif (data not shown), whereas PTS rate at HF_slip_eGFP was 14.37% for SNI and 0.87% for SND (Figure 7C). Interestingly, these are not as high as slippage rates when using only the genome as template (Figure 7B), indicating that the accumulation of subgenomic RNA species with +/-A at the slippage motif is somehow constrained. As no difference in the transcription rate of diverse subgenomic RNA variants is anticipated, then this result fits with two putative scenarios: (i) PVX polymerase is more prone to slip when replicating the viral genome than during transcription of the subgenomic RNA, or (ii) the stability of subgenomic RNA variants with +/-A at the slippage motif is lower than the variant without insertion/deletion. The second scenario is not the most likely as, indeed, the RNA variant with no insertion/deletion has a stop codon in the main frame just downstream the slippage motif; therefore, it is not translated (not protected/stabilized by ribosomes) and has a longer 3’ UTR (more prone to be recognized and degraded by the NMD mechanism).

Although not conserved in all reported strains, most PVX genomes annotated at NCBI, including the one used in this study, harbour an A_6_ run at the viral replicase coding sequence, specifically at the 3’ end of the segment that codes for the methyltransferase domain (Figure 7A). We used the above RNA samples as template to generate RT-PCR amplicons spanning this motif (Figure 7A) and, by analysing the NGS data, found that PTS events take place at this position, although at lower level than at the HF_slip site (Figure 7C). In this particular case, the so-produced viral RNA species with +/-A at the slippage motif do not give access to an overlapping ORF; instead, they produce truncated versions of the viral replicase including only its methyltransferase domain. In fact, truncated fragments of the PVX polymerase have been detected during infection (46). Therefore, PTS may play a role here in the production of a free methyltransferase, thus mimicking what other viruses do via different strategies (e.g., nsP1 from alphaviruses (47)).

## Discussion

Even though the conservation of genome sequences just upstream of the PIPO coding sequence had already anticipated that *pipo* ORF was acceded via PTS in all viruses from the family *Potyviridae*, empirical demonstration was just available for a few members of the *Potyvirus* genus. Our work, along with other recently published studies, indicate that PTS indeed takes place not only in potyvirids from other genera, but also in a plant RNA virus from another family (48, 49, and this work). Similar observations in animal-infecting viruses, in both naturally-occurring and artificially-introduced slippage motifs at Ebolavirus genome (7, 12) and Theiler’s murine encephalomyelitis virus genome (50), respectively, have been reported. Together, these results support the idea that viral polymerase slippage is a widespread feature of RNA viruses in general.

Notably, the data depicted in Figures 1 and 3 unveil a previously underestimated phenomenon of PTS: the production, during infection, of viral RNA species harbouring a significant proportion of SND at slippage motifs, comparable in frequency to those with SNI. Although SND events do not provide access to overlapping ORFs in CocMoV, the elevated frequency of these particular molecules compels us to hypothesize potential roles and evolutionary implications for the production of RNA species coding for truncated proteins. Regarding roles, two non-mutually exclusive scenarios are plausible: (i) they work as a negative self-regulatory strategy to avoid viral overaccumulation and the deleterious consequences of virus diseases for the host, as truncated proteins might not be functional; (ii) they are translated into truncated, yet functional, proteins so enlarging the coding capacity of viral genomes. Indeed, an example that supports the feasibility for this second scenario has been reported for ClYVV (13), and it might be also the case reported here of the sole MTS domain expressed during PVX infection (Figure 7D). An alternative option within this second scenario is exemplified by the case of ANSSV, where polymerase slippage results in RNA species with premature stop codons in close proximity to sequences encoding protease cleavage sites, so having the potential to complement the action of viral proteases (Figure 5).

In terms of evolution, the production of RNAs coding for truncated proteins fits well with a parsimonious evolutionary trajectory that potentially explain the appearance of *pipo* ORF in all potyvirids (Figure 8). It seems logical to assume that the genome of a common ancestor for all potyvirids encoded from P3 to CP, with P3 and CI playing roles in cell-to-cell movement (51). The next step would have included the appearance of a repetition of As in the central region of P3, which promoted PTS (SNI/SND) and, consequently, the appearance of few viral RNA species with a premature stop codon. These peculiar RNAs, when translated, would have produced a truncated form of P3, which took over the role of P3 during virus cell-to-cell movement, so resembling P3N-ALT from ClYVV (13). In that scenario, the hypothesized role of P3 in virus movement would have not been necessary, meaning that the 3’ half of P3 coding sequence could evolve to either improve another former P3 role or acquire a new one. This idea, in fact, is in line with recent analyses showing that the region of potyviral genomes coding for the C-terminal part of P3, but not that coding for the N-terminal part of it, displays high variability (52). Later during evolution, from PTS events induced by the homopolymeric run of As, the SNI would have been able to explore an alternative frame in order to extend the size, and improve the role, of P3N-ALT in virus cell-to-cell movement, so giving rise to P3N-PIPO. After different genera evolved, some viruses seem to have specialized to just introduce an A at the slippage motif to produce P3N-PIPO, whereas others seem to still produce P3N-ALT and P3N-PIPO at different ratios, depending on their ability to delete/add a single A at the slippage motif during replication. Further evolution events had to be necessary to explain the appearance of a U_8_ slippage motif (and the disappearance of A_6_) upstream of the *pipo* ORF in WMVBV (Figure 4).

**Figure 8.**
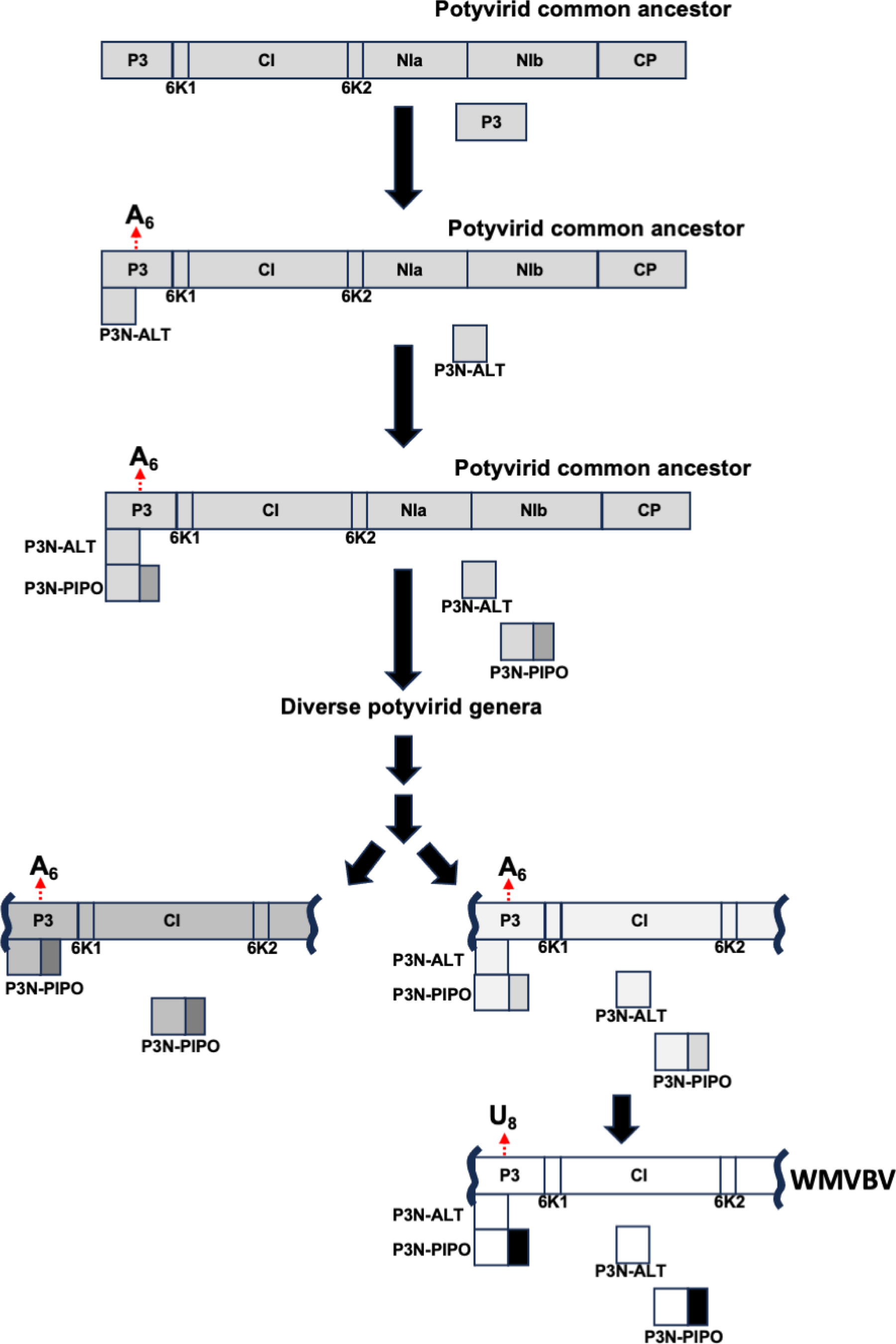
Hypothetical evolutionary trajectory of potyvirids that explains the appearance of a slippage motif and the *pipo* ORFs in the P3 coding sequence. Schematic representation of the coding sequences from ancestral potyvirids, as well as P3-related factors associated to potyvirid cell-to-cell movement, at each step in the evolution of the *Potyviridae* family. See the main text for more information.

A striking conclusion from our analysis of all available potyvirid full-length genome sequences (Supplementary Table S1) is that, apart from the conserved slippage motifs located upstream of *pipo* and *pispo* ORFs in all potyvirids and in a subset of potyviruses infecting sweet potato, respectively, many specie-specific A_n_/U_n_ (n ≥ 6) motifs are present in certain viral genomes. Importantly, some of them would give access to short ORFs with more than 30 amino acids (e.g., catharanthus mosaic virus, cucurbit vein banding virus), while the others would promote the expression of truncated versions of viral proteins. Having in mind that homopolymeric runs of nucleotides formed by A_6_, A_7_, U_6_ or U_7_ are underrepresented in potyvirid genomes (8, 9), it is then reasonable to propose that these motifs were retained during evolution since PTS at those positions play a role during the course of viral infections. On the other hand, the analysis shown in Figure 4 with ANSSV suggests that not all A_n_/U_n_ (n ≥ 6) motifs, when comparing to other sequences, seem to induce the production of RNA molecules having SNI/SND. A previous study with Ebola virus supports this idea, as it showed that the viral replicase slips over an A_7_ motif to introduce an additional A in the GP coding sequence, but not in an equivalent motif in the sequence coding for the L protein. In that report, authors demonstrated that a short loop located just upstream of the slippage motif is critical for this SNI to happen (12). In the case of potyvirids, a report based on TuMV indicated that nucleotide sequences flanking the homopolymeric run of As is also relevant; however, in that particular case, it is the GC content rather than the RNA secondary structure that influences PTS (11). Moreover, as an additional layer of complexity, we also found that factors unrelated to cognate viruses, such as the coinfection with other viruses and the identity of infected hosts, also influence PTS rates (Figure 5 and Supplementary Figure S1). The fact that different hosts do not accumulate the same proportion of slippage products might be due to their intrinsic capacity to prevent the accumulation of RNAs harboring premature stop codons with large 3’UTRs, such as those produced by PTS. Indeed, we found strong evidences that NMD is, at least in part, preventing the accumulation of natural slippage products in *N. benthamiana* plants infected with PPV (Figure 5B). Importantly, when we were preparing this manuscript, results from an independent group showed a comparable impact of NMD over the rate of slippage products in TuMV by using a different approach (53). Intriguingly, the impact of NMD on potyvirid infections remains a subject of discussion due to conflicting results (53–55).

As demonstrated in this study and others, numerous factors have the potential to influence PTS rates either positively or negatively, so that it appears necessary to study this phenomenon on a case-by-case basis. Nevertheless, given the impact that genuine PTS has over the viral proteome, it will be exciting in the future to identify new examples of protein expression via this strategy in RNA viruses, and investigate not only the reasons and consequences of PTS alterations induced by diverse factors, but also the mechanisms promoting these changes.

## MATERIALS AND METHODS

### Plants, virus isolates and bacteria strains

Melon plants (cv. Védrantais) were grown in a greenhouse with 16h/8h light/dark cycles at 22-to-25°C. *Nicotiana benthamiana*, *Ipomoea nil* and *Cucurbita pepo* (cv. Diamant F1) plants were grown in a greenhouse at 24-to-28°C (day) and 22-to-24°C (night) with a natural photoperiod (no less than 12 h of light, supplemented with artificial illumination if needed). The *Areca catechu L.* tree was grown in the field.

Natural isolates of coccinia mottle virus (CocMoV isolate Su12-25, *Ipomovirus* genus, *Potyviridae* family, GenBank accession no. KU935732) (24), wild melon vein banding virus (WMVB isolate Su03-07, *Potyvirus* genus, *Potyviridae* family, GenBank accession no. KY623506) (25), cucurbit yellow stunting disorder virus (CYSDV isolate AILM, *Crinivirus* genus, *Closteroviridae* family, GenBank accession no. AY242077 and AY242078) (26) were maintained in melon plants. A natural isolate of cucumber vein yellowing virus (CVYV isolate AILM, *Ipomovirus* genus, *Potyviridae* family, GenBank accession no. MK777994) (27) was maintained in *Cucurbita pepo*. A natural isolate of sweet potato virus 2 (SPV2 isolate AM-MB2, *Potyvirus* genus, *Potyviridae* family, GenBank accession no. KU511270) (23) was maintained in sweet potato plants. A natural isolate of areca palm necrotic spindle-spot virus (ANSSV isolate HNBT, *Arepavirus* genus, *Potyviridae* family, GenBank accession no. MH330686) (28) was identified and maintained in an *A. catechu L.* tree in the field (Hainan, China).

A previously built infectious clone of watermelon mosaic virus (WMV, *Potyvirus* genus, *Potyviridae* family) (29) was used to inoculate melon plants in order to get fresh infected tissue for further manual inoculations.

*Agrobacterium tumefaciens* strains carrying either pBIN-UPF1, pBIN-U1D or pBIN-P14 (30) were kindly provided by Prof. Daniel Silhavy (Institute of Plant Biology, Eötvös Loránd Research Network, Szeged, Hungary).

### Primers

A list of all oligonucleotides/primers used for this study, with their corresponding sequences, is shown in Supplementary Table S2.

### Plasmids

Plum pox virus-(PPV-) derived constructs with an out-of-frame eGFP preceded by the indicated slippage motifs were built by LR recombination by using the previously described pGWBin-PPV (31) as destination vector, and the corresponding pDONR-slippage_motif_eGFP variants as entry clones. These were prepared by BP recombination between pDONR207 (Invitrogen) and synthetic DNA strings harbouring (from 5’ to 3’): attB1, the corresponding 21-nt slippage motifs, the eGFP coding sequence with synonymous mutations avoiding stop codons in the +2 frame (+1 frame once this fragment is inserted in the virus after BP and LR), and attB2 (Supplementary File S1).

The infectious cDNA clone of CocMoV was built by using a recombination method in yeast, as previously described (29). A detailed description of the methodology is included in the Supplementary File S2, and the sequence of the CocMoV full-length clone, termed pCocMoV_Su12-25r, was deposited in NCBI (GenBank accession no. OL744323). For the introduction of an extra PTS motif in CocMoV, we modified the pCocMoV_Su12-25r plasmid by taking advantage of its unique *Spe*I restriction site located at the CocMoV P1 coding sequence. Inserts were first generated as small double-stranded DNAs by *in vitro* annealing of specific oligonucleotide pairs (#1/#2 and #3/#4), and then introduced in the *SpeI*-linearized plasmid by ligation to get the corresponding pCocMoV_Su12-25r derivatives.

The PVX-derived constructs pGWC-PVX-eGFP, pGWC-PVX-HF_slip_eGFP and pGWC-PVX-HF_slip_mut_eGFP were built by LR recombination, using the previously described pGWC-PVX (32) as destination vector, and pENTR1A-eGFP, pENTR1A-HF_slip_eGFP and pENTR1A-HF_slip_mut_eGFP as entry clones, respectively. The constructs were prepared by PCR amplification of eGFP from pLONG-GFP (33) with primer pairs #5/#8, #6/#8 and #7/#8 to generate ATG_eGFP_stop ATG_HF_slip_eGFP_stop and ATG_HF_slip_mut_eGFP_stop fragments, respectively, and further blunt-end ligation of these products into pENTR1A previously digested with *Xmn*I and *EcoR*V.

### Virus inoculation

For manual inoculation, 15 µl of crude extract from systemically infected plants (1 g in 2 ml of 5 mM sodium phosphate, pH 7.2) were finger-rubbed on young leaves of 1-month-old plants previously dusted with carborundum.

For inoculation of WMV-, CocMoV- and PPV-derived full-length clones, biolistic with HandGun (34) for the first two clones and Helios Gene Gun System (Bio-Rad) (35) for the last clone was done as previously described.

### Transient expression of proteins

Co-expression of either UPF1 or U1D along with the silencing suppressor P14 from pothos latent virus in upper non-inoculated leaves of PPV-infected *N. benthamiana* plants was carried by infiltration of *A. tumefaciens* strains harboring the corresponding plamids (30). To do that, we prepared cell cultures and the indicated mixes of agrobacteria as previously described (36).

### RNA extraction, next-generation sequencing, and data analyses

Upper non-inoculated leaves of plants infected with the indicated viruses were used to extract total RNA with TRI Reagent® (Sigma), as indicated by the manufacturer. Then, 1 μg of total RNA was used to generate either amplicons or cDNA libraries for next-generation sequencing.

To generate amplicons, RNA samples (from three independent plants in all cases) were subjected to reverse transcription with the AMV enzyme (Promega) and random hexanucleotides as primers, and the so-generated cDNAs were then used as template for the PCR amplification of 250-to-300-nt fragments spanning the indicated slippage motifs. Primer pairs #9/#10 and #11/#12 were used to get amplicons spanning the PIPO and the artificially-introduced slippage motifs in PPV, respectively. Primer pairs #13/#14, #15/#16 and #17/#16 were used to get amplicons spanning the naturally-occurring, as well as a long and a short fragment of artificially-introduced slippage motifs in PVX, respectively. Primer pairs #18/#19, #20/#21 and #22/#23 were used to get amplicons spanning PIPO, P1-ALT and the artificially-introduced slippage motifs in CocMoV, respectively. Primer pairs #24/#25 and #26/#27 were used to get amplicons spanning PIPO and PISPO in sweet potato virus 2 (SPV2), respectively. Finally, primers #28/#29, #30/#31 and #32/#33 were used to get amplicons spanning PIPO in watermelon mosaic virus WMV, CVYV and WMVBV, respectively. These RT-PCR products were sent to the genomics facility at Madrid Science Park. Pair-end sequencing (2x300) was done with MiSeq Reagent Kit v3 in a MiSeq platform (Illumina) by following the manufacturer’s instructions.

To analyse the transcriptome of an areca palm leaf infected with ANSSV, total RNA was sent to Biowefind Biotechnology Co., Ltd. (Wuhan, China). After rRNA depletion, RNA was used to prepare the library with NEBNext Ultra Directional RNA Library Prep Kit, which was further sequenced with a Novaseq 6000 platform (Illumina) by following the manufacturer’s instructions.

Sequencing reads were aligned against the genome sequence of their corresponding viruses with bowtie2 (37). These alignments were analysed with SAMtools mpileup (38) to further generate lists of single-nucleotide polymorphisms, as well as small insertions/deletions with their corresponding frequencies at each position, by using in-house Perl scripts (available upon request). Statistical differences between PTS rates were tested with the post-hoc Tukey HDS test.

### Fluorescence imaging

GFP fluorescence in infected plants was observed with an epifluorescence stereomicroscope using excitation and barrier filters at 470/40 nm and 525/50 nm, respectively, and photographed with an Olympus DP70 digital camera.

## ACKNOWLEDGMENTS

We are grateful to Dániel Silhavy for providing *A. tumefaciens* strains carrying pBin61-P14, pBin61-UPF1 and pBin61-U1D. This work was supported by grant BIO2015-73900-JIN funded by AEI-FEDER to AAV, as well as PID2019-105692RB-100 and PID2019-110979RB-I00 to JJLM and AAV, respectively, funded by MCIN/AEI/10.13039/501100011033. The Center for Research in Agricultural Genomics (CRAG) was also supported by grants SEV-2015-0533 and CEX2019-000902-S funded by MCIN/AEI/10.13039/501100011033, and by the CERCA Programme, Generalitat de Catalunya. MLD was supported during her work at CRAG and at INRA-Avignon by FPI contract BES-2014-068970 from MICIN and FEDER. Funders had no role in study design, data collection and interpretation.

---

**Supplementary Figure S1. The effect of co-infections on CocMoV PTS rates.** Slippage frequencies at the GA_6_ motif in *alt* (upper panel) and at the GA_7_ motif located upstream of the *pipo* ORF in CocMoV (bottom panel). Bars represent the average frequencies (indicated between brackets) for single nucleotide insertions (SNI) and single nucleotide deletions (SND) at the indicated slippage motifs. Error bars represent standard deviation (n = 3). Statistical differences were tested with the post-hoc Tukey HDS test (* *p* value < 0.05; ** *p* value < 0.01).

**Supplementary Figure S2. The conservation of the U_8_ motif in isolates of WMVBV**. Alignment of a segment from the P3 coding sequence from Su12-21 and the genome sequence from Su03-07, two WMVBV isolates. The presence of a conserved U_8_ motif is highlighted by a red square. Two specific nucleotide differences between these isolates are indicated by red asterisks.

